# What is hidden in the darkness? Characterization of AlphaFold structural space

**DOI:** 10.1101/2022.10.11.511548

**Authors:** Janani Durairaj, Joana Pereira, Mehmet Akdel, Torsten Schwede

## Abstract

The recent public release of the latest version of the AlphaFold database has given us access to over 200 million predicted protein structures. We use a “shape-mer” approach, a structural fragmentation method analogous to sequence *k*-mers, to describe these structures and look for novelties - both in terms of proteins with rare or novel structural composition and possible functional annotation of under-studied proteins. Data and code will be made available at https://github.com/TurtleTools/afdb-shapemer-darkness

## 1 Introduction

The universe of all possible protein-like sequences is sparsely populated by natural proteins (Levitt, 2009) and, with the contribution of both experimental and computational approaches, has been both greatly expanded and functionally annotated. Recent deep-learning based protein structure prediction techniques, such as AlphaFold (Jumper *et al*., 2021), are now also expanding our knowledge at the structural level. Apart from the hundreds of thousands of experimentally resolved structures in the Protein Data Bank (PDB) (Berman *et al*., 2000), 214 million protein structural models are publicly available since July 28 2022 through the AlphaFold database version three (AFDB v3) (Varadi *et al*., 2022; Callaway, 2022). These structures open new avenues for the better understanding of the diversity of folds adopted by all natural proteins and their potential biological roles. However, this data explosion imposes the need for efficient and automatic methods for protein structure analysis and comparison.

Here, we decompose structures into fragment shape-mers analogous to *k*-mers in a protein sequence, or words in natural language. This allows for fast and efficient large-scale comparison across millions of structures using natural language processing (NLP) techniques. With this framework in place, we attempt to describe the structural novelties in AFDB v3 compared to the PDB with two different perspectives in mind - first, differentiate between the distributions of PDB and AFDB structural folds and pinpoint structural outliers in AFDB; and second, connect unannotated and functionally unknown proteins to similar structures for which some functional annotations are available.

Two recent efforts in similar directions include van Kempen *et al*. (2022), where a 20-letter structural alphabet was created based on the local distances and angles between each residue and its closest neighbor in order to use fast sequence search approaches, and Bordin *et al*. (2022), where the authors combine sequence profile searches, language models, and structure searches to assign parts of AFDB v1 structures to CATH domains (Orengo *et al*., 1997). Unlike the former, our approach allows for the understanding of larger fragments encompassing whole secondary structure elements and their structural context. And, complementary to the latter, it allows for purely structural comparisons across millions of structures in a completely unsupervised fashion, thus bringing to light not only structural outliers in terms of known domains, but also outliers in terms of structural fragment content.

## 2 Methods

### 2.1 Shape-mer construction

#### Rotation-invariant moments

The shape-mer approach used builds upon the Geometricus algorithm from Durairaj *et al*. (2020), which used 4 rotation invariant moments calculated on fragments of protein structures, defined either using overlapping *k*-mers of backbone *α*-carbon coordinates, or *α*-carbon coordinates within a sphere of a certain radius of each residue. Here, we extend our approach to make use of 17 rotation invariant moments including 3 second order moments from Mamistvalov (1998), 12 third order moments from Flusser *et al*. (2003, 2016) and the chiral invariant moment from Hattne and Lamzin (2011). These 17 moments were calculated for 4 different fragmentation types on *α*-carbon coordinates: *k*-mers of size 8 and 16, and spheres of radii 5 and 10; for a total of 68 moments for each residue in a protein, ignoring the first 8 and last 8 residues.

#### Neural network

We trained a neural network using PyTorch (Paszke *et al*., 2019) with input size 68 (i.e. the number of moment invariants calculated per residue), 1 hidden layer of size 1024 and output size 10. The network was trained with contrastive loss to reduce the output distance between structurally equivalent pairs of residues and increase the distance between non-equivalent pairs in the training set. As the output of the network is 10 floating point numbers between -1 and 1, this could be discretized into 10 bits based on whether the value was greater than or less than 0, resulting in 1024 shape-mers, where the bitwise Hamming distance defines the similarity between shape-mers. See Figure 4 for a comparison of these extended shape-mers against Geometricus and TM-score.

#### Training data

To train shape-mer representations from the moment invariants, we used structures from the CATH database having <40% sequence identity that could be assigned CATH functional family (funfam) assignments with an E-value < 10*e*6−10. From these 8333 structures, US-align (Zhang *et al*., 2022) was used to align and superpose all pairs within each funfam cluster and 3 * 8333 randomly chosen pairs across clusters. Pairs of residues from two proteins belonging to the same funfam, with an alignment TM-score > 0.8 and a weighted RMSD of the 16 residue sequence neighborhood and 10 Åstructure neighborhood < 2 were considered as positive pairs. Pairs of residues from two proteins belonging to different CATH superfamilies, with an alignment TM-score < 0.6 and a weighted neighborhood RMSD > 5 were considered as negative pairs. This resulted in a total of 3,398,839 residue pairs for training. Training took 30 mins on 1 RTX-3080TI.

#### Shape-mer calculation

Shape-mers were calculated for all AFDB v3 proteins (214M proteins modelled as monomers) and every chain of the PDB (version PDB_220907T050415, 652,611 chains). For AlphaFold models, we followed the approach described in the analysis of AFDB v1 (Akdel *et al*., 2021) to split each protein into segments with Gaussian smoothed plDDT > 70. Shape-mers were then calculated for each segment and concatenated. Computations took about 1 week using 20-40 threads in our cluster. Proteins with less than 20 amino acids were ignored. Figure 1 shows four example shape-mers along with corresponding examples of proteins with over 50% of their structure made up of that shape-mer.

**Figure 1:**
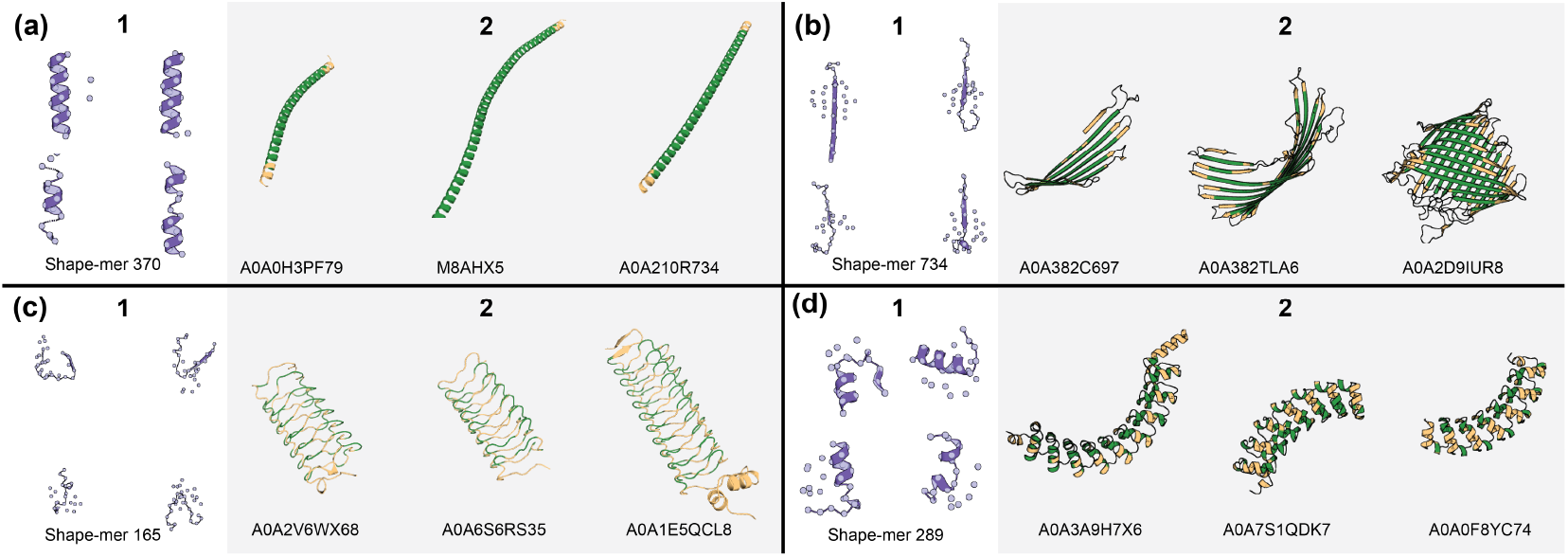
Examples of shape-mers and proteins containing them. Residues associated with the corresponding shape-mer are colored in green.

### 2.2 Natural language processing techniques

As shape-mers within a protein structure can be considered equivalent to words within a document, we use two NLP techniques to compare shape-mers of protein structures. We train Word2Vec (Mikolov *et al*., 2013) on shape-mers from the PDB dataset using gensim (v4.2.0) (Rehurek and Sojka, 2011) with a window size of 16 and embedding size of 1024. Protein-level embeddings are obtained by averaging across normalized shape-mer embeddings. Non-negative Matrix Factorization (Lee and Seung, 2000) as implemented in scikit-learn (v1.1.1) (Pedregosa *et al*., 2011) was trained with 1000 topics on the term frequency inverse document frequency matrix of the shape-mers for all PDB chain structures. Each topic is a weighted combination of the 1024 shape-mers and each protein is a weighted combination of these topics. We assigned proteins to each topic using knee detection as implemented in the kneed Python library (v0.7.0) (Satopaa *et al*., 2011) with a weight cut-off *S* of 4. Topic-specific residue scores were obtained by multiplying the corresponding topic weight for each shape-mer with the normalized Euclidean distance between each residue and the central residue of the shape-mer, aggregated across all shape-mers.

### 2.3 AlphaFold database version 3 analysis

#### Dataset collection

We mapped each entry in AFDB v3 to its corresponding UniRef50 cluster (v03.08.2022) (Suzek *et al*., 2007) and computed the distribution of average pLDDTs across all elements in each cluster. Given the diversity of pLDDTs for entries within the same UniRef50 cluster (Figure 2), we choose as representative of each UniRef50 cluster the longest entry with an average pLDDT above 70, and restricted our study to those with an average pLDDT above 95. This resulted in total to 1’066’673 structures, referred to as UniRef50-95. The most common type of repeat protein within the dataset corresponds to 21,300 coiled coil-like long helical repeats (shown in Figure 1A) consisting almost entirely (>50%) of shape-mer 370 - these were removed from further analyses.

**Figure 2:**
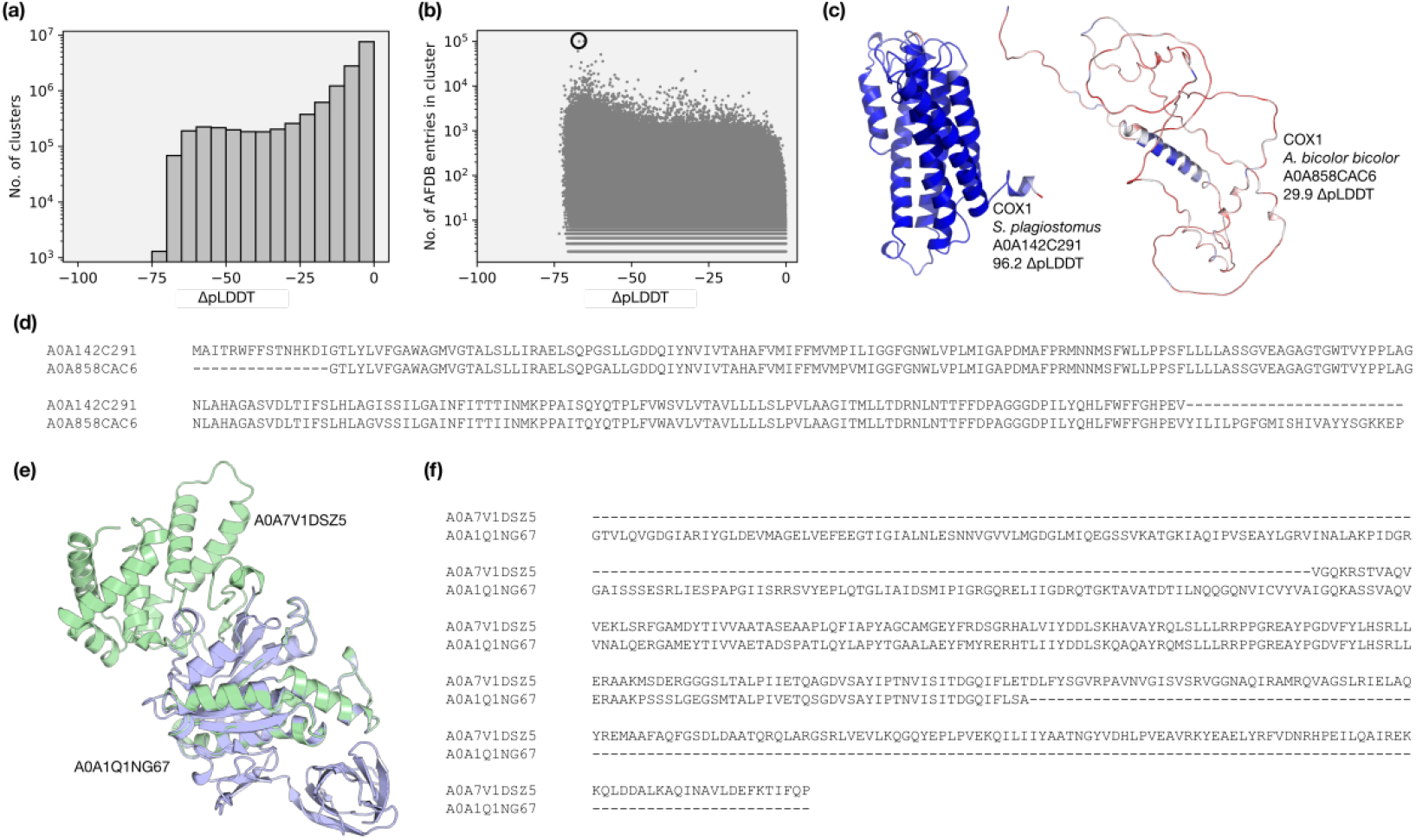
plDDT analysis of UniRef50 **(a)** Distribution of ΔplDDTs (difference between lowest and highest average plDDT) across all clusters **(b)** Distribution of ΔplDDTs vs. cluster size **(c)** Highest and lowest confidence models in the most populated cluster and **(d)** corresponding sequence alignment. **(e)** Two high confidence proteins from UniRef50_P56757 with differing structures and **(f)** corresponding sequence alignment.

#### Novelty and outlier detection

We use the Isolation Forest algorithm (Liu *et al*., 2008) from scikit-learn to detect outliers in AFDB as compared to the PDB. The forest was trained on the Word2Vec embeddings of the PDB dataset with 5% contamination rate and used to detect outliers in the UniRef50-95 dataset. To obtain similarities between proteins, an approximate *k* nearest neighbors graph was created using PyNNDescent (v0.5.7, Euclidean metric and 30 neighbors) (Dong *et al*., 2011) across both the PDB and UniRef50-95 dataset. For sets of proteins of interest, graphs connecting each protein to up to three nearest neighbors with Word2Vec embedding distance < 0.15 (see Figure 6) are constructed to perform connected components analysis analysis with NetworkX (v2.8.6) (Hagberg *et al*., 2008).

#### Functional brightness and darkness assignment

We defined the functional “brightness” of a given protein as the maximum full-length coverage with family, domains and predicted structural features (coiled coils and predicted intrinsic disorder) of the best annotated entry in its corresponding UniRef50 cluster. These annotations were extracted from UniProt, UniParc and Interpro, (uni, 2021; Blum *et al*., 2021) excluding those containing ‘DUF’, ‘Hypothetical’, ‘Putative’ and ‘Uncharacterised’ in their descriptions. The higher the coverage (i.e., the closer to 100%) of the best annotated element in a UniRef50 cluster, the brighter are all of its corresponding entries. Similarly, the lower the coverage (i.e., the closer to 0%) of the best annotated element in a UniRef50 cluster, the darker are all of its corresponding entries. We considered as “dark” proteins those where the brightest UniRef50 representative has a brightness lower than 5%. Figure 5 highlights the distribution of brightness across UniRef50 and AFDB v3 at different pLDDT levels.

## 3 Results

### 3.1 AlphaFold modelling differs within the same UniRef50 cluster

Driven by multiple independent observations that highly similar protein sequences were modelled in AFDB at different levels of predicted accuracy, we analysed how widespread these cases are within UniRef50 clusters. We observed that for 53% of all UniRef50 clusters with more than one element in AFDB the difference in pLDDT between the lowest and the highest confident models (ΔplDDT) is just 5 pLDDT units, but at least 10% of the clusters encompass models where the absolute ΔplDDT is larger than 30, with the largest being 70 (Figure 2a).

This variation does not correlate with the size of the UniRef50 cluster (Figure 2b) neither is affected by the presence of fragments. One example is given in Figure 2c-d. The largest cluster in the dataset is UniRef50_Q9MIY8 and encompasses members of the Cytochrome C oxidase subunit 1 (COX1) protein family, with more than 100K entries in AFDB. The absolute ΔpLDDT is larger than 60, but the highest and lowest confidence models have identical sequences.

To confirm that high confidence models from similar sequences have similar modelled structures, we performed a Word2Vec connected component analysis on all structures with over 95 plDDT from the 100 most populated UniRef50 clusters. For each UniRef50 cluster, most proteins (>99%) with length within one standard deviation were in the same connected component. Smaller proteins (< 100 residues), which are likely fragments, formed a number of smaller disconnected components. To ensure that we only analyse robust, high quality models, we continue with the UniRef50-95 dataset (described in Methods). While on average these represent the structures seen in the entire cluster, Figure 2e-f shows an example where this is not the case.

### 3.2 What’s new in AFDB predicted models across UniRef50

Ever since the first version of AFDB was made available in the summer of 2021, various efforts focused on the identification of structural novelties in the thousands of available predicted models (Akdel *et al*., 2021; Bordin *et al*., 2022). Our shape-mer approach combined with classic NLP techniques allows easy scaling of such analyses to millions of structures in a matter of days. In addition, the methods described inspect distributions of structural content in an unsupervised and alignment-free manner, thus providing a different perspective from alignment-based and annotated fold-based methods.

The difference between the frequency of different topics in the PDB and the AFDB highlights the types of structural motifs that have been now expanded by AFDB at an unexpected rate. For example, the frequency of the topic describing transmembrane regions (Figure 7A) in AFDB is 8% higher than in the PDB. Similarly, outlier detection highlights those structures in AFDB that are less common in PDB and thus may represent novel folds or quirks of AlphaFold that may need to be considered when interpreting AlphaFold models (Figure 8).

One important application of our method is for the analysis of poorly annotated and functionally dark proteins. Figure 3a shows a set of mostly functionally dark proteins that share a common fold and cluster closely with known enzymes that bind amino acids (mainly arginine) and are involved in their metabolism. This suggests that these dark proteins may adopt the same fold and, thus, carry a similar function. In Figure 3b, we show a cluster containing Outer Membrane *β*-Barrels but also other structurally similar proteins. In the middle of this cluster, there are a number of structural outliers (bigger circles) that consist of single layer beta-sheets, which are rare amongst natural proteins (Xu *et al*., 2019). Many of these are Gloverins, antimicrobials targetting Gram-negative bacteria for which no homologs exist in the PDB (Yi *et al*., 2014). Finally, Figure 3c shows a cluster of mostly dark and structural outliers that cluster together with adhesin-like and pectin lyase domains. While a few of these have been experimentally characterized, AFDB indicates that many more are to be annotated.

**Figure 3:**
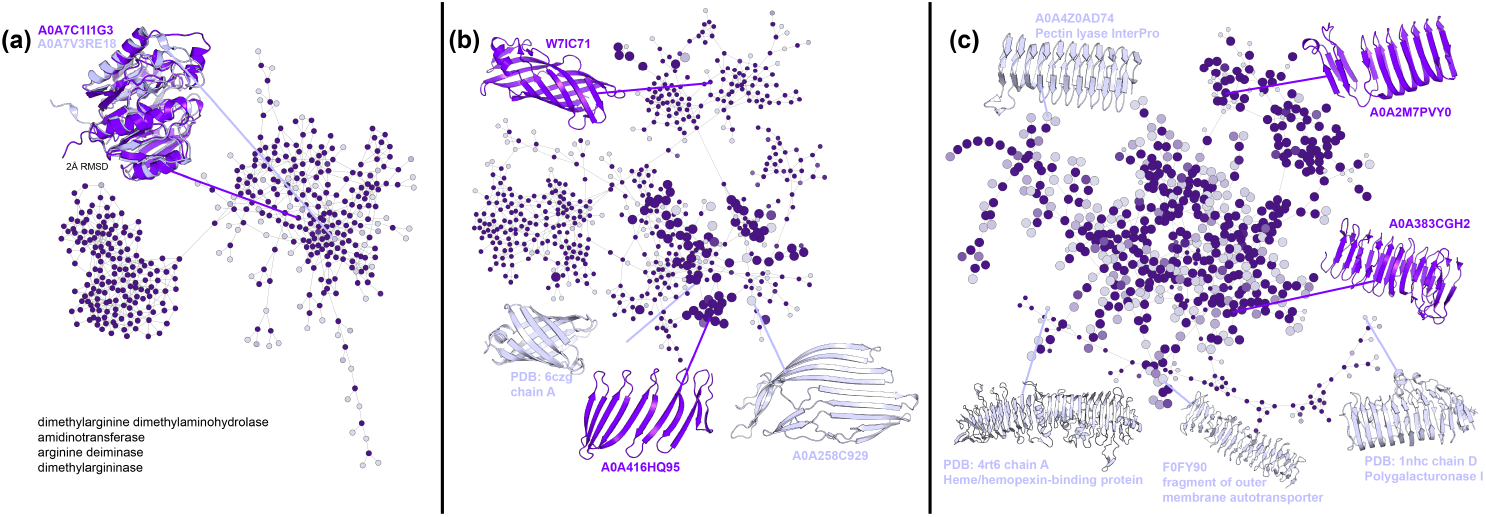
Examples of functionally dark proteins and their closest UniRef50-95 and PDB Word2Vec neighbors. Purple circles denote dark proteins, and grey circles represent functionally or experimentally characterized proteins. Large circles mark structural outliers.

As average Word2Vec embeddings lose information about context beyond what is stored in the individual word (shape-mer) vectors, structures with similar composition but different orders are seen as similar. While this may be an interesting way to explore phenomena such as circular permutations, a powerful next step is to train a language model on the sequences of shape-mers to truly capture structural context.

## 4 Conclusion

AFDB v3 has brought forth a sudden and dramatic structural coverage across the tree of life, and is extensively used by life scientists. However, the confidence of the predicted models vary widely even across very similar sequences, conveying a false sense of structural diversity. This problem has been identified by the AFDB maintainers and is now being rectified as part of the version 4 release, as stated in the AFDB FAQ.

Still, our structural fragment-based analysis shows that AFDB v3 indeed expands structural diversity even when considering the most confident of robust predictions. Some fold spaces have been considerably expanded, and a number of models are distinct from experimental structures. A substantial number of unannotated and under studied proteins are modelled with high confidence and many are found to be structurally identical to proteins with functional annotations from various manually and automatically curated databases, opening the doors to automated annotation transfer.

Such new ways of defining discrete structural fragments with spatial context, and no underlying fragment database, have an edge in two different ways. Firstly, they give us access to the myriad of hyper-optimized sequence-based techniques (search, alignment, motifs, profiles, trees and more). Secondly, they are more analogous to words in natural languages, thus making them better suited to the recent NLP approaches that are revolutionizing a variety of fields.

### Broader Impact

Researchers in bioinformatics and life sciences in general may benefit from such a large-scale overview of the differences and similarities between experimental and computationally resolved structures. Our shape-mers or structural words enable easy navigation through the protein universe from a fragment perspective, also potentially useful for protein engineering and design techniques. Given the increasing success of language models in a variety of different tasks, the recent explosion of structures can be combined with the ideas proposed here to create protein embeddings combining both structure and sequence information across the protein universe.

## A Appendix

**Figure 4:**
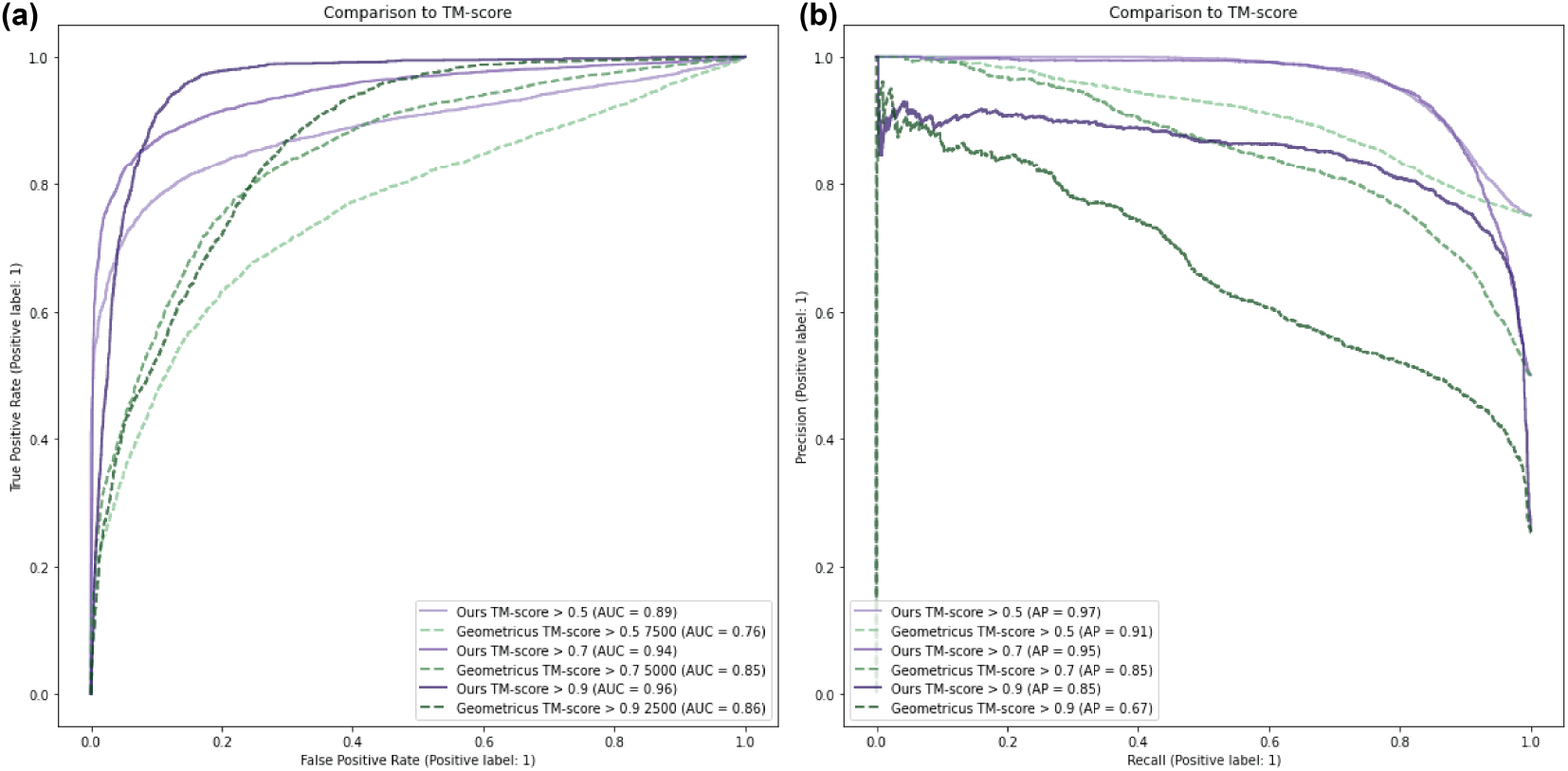
10,000 pairs of proteins from SCOP (with <40% sequence identity from each other) were gathered such that 2,500 pairs each have TM-score <0.5, 0.5-0.7, 0.7-0.9 and >0.9. Dynamic programming alignment was performed on the Geometricus shape-mers and our extended and trained shape-mers across these pairs. Panels **(a)** and **(b)** show the ROC and Precision-Recall curves at different TM-score thresholds.

**Figure 5:**
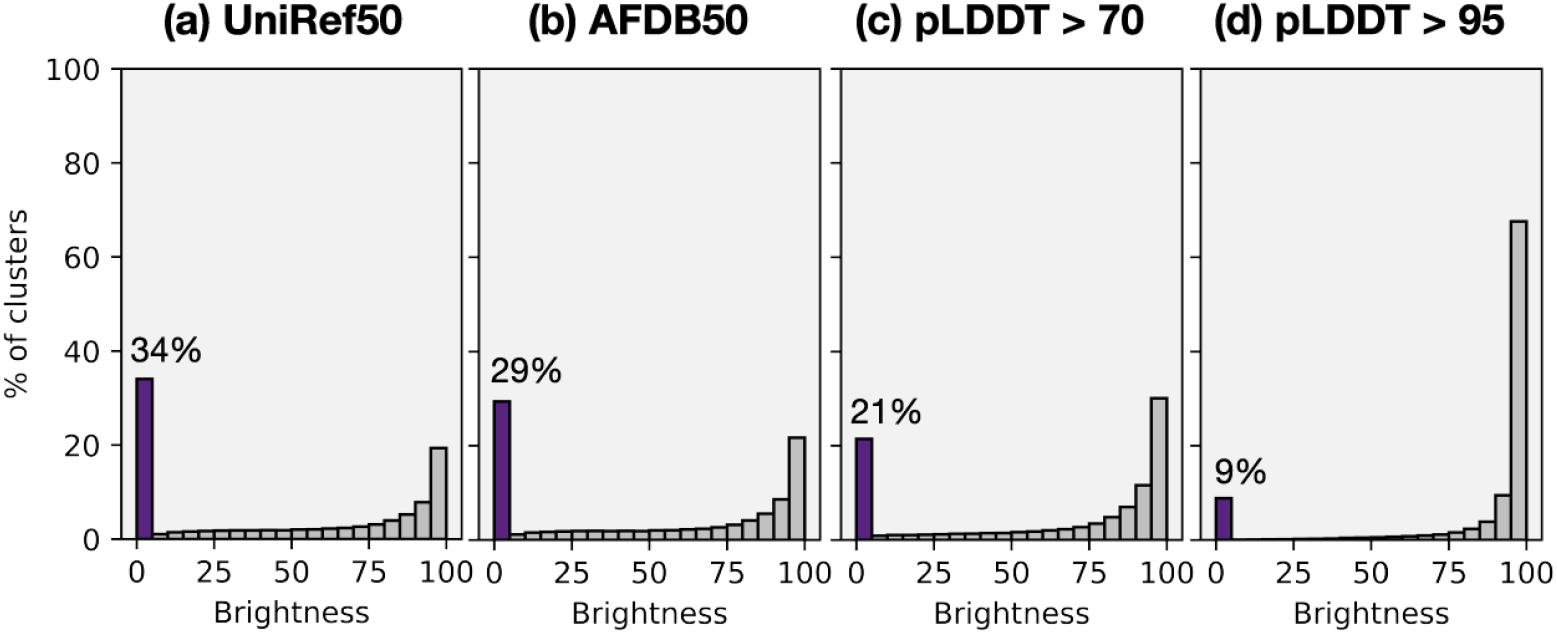
Functional brightness in UniRef50 and AFDB **(a)** Distribution of annotation levels across (1) UniRef50 (2) UniRef50 clusters with AFDB representatives (3) with plDDT > 70 and (4) with plDDT > 95. Data collection and functional brightness calculations took, in total, about 2 weeks on a single CPU.

**Figure 6:**
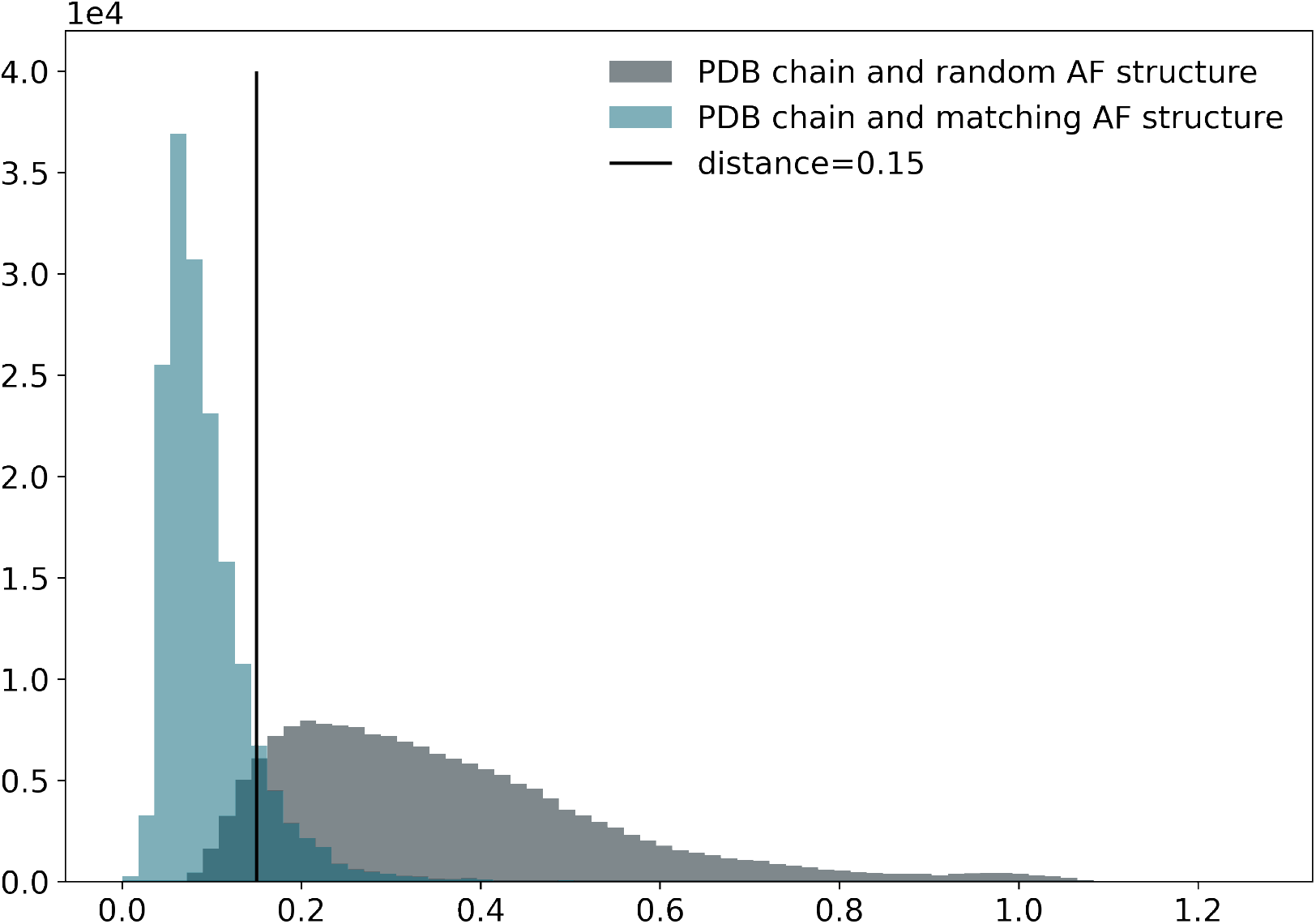
Euclidean distances between Word2Vec shape-mer embeddings were calculated for 168,010 PDB chains and their corresponding UniProt AFDB models (mapping data obtained from SIFTS) and compared to distances between the same PDB chains and randomly selected SwissProt AFDB models. A threshold of 0.15 distance is chosen from this plot and used in the manuscript as 89% of matching pairs lie under this threshold compared to 7% of random pairs.

**Figure 7:**
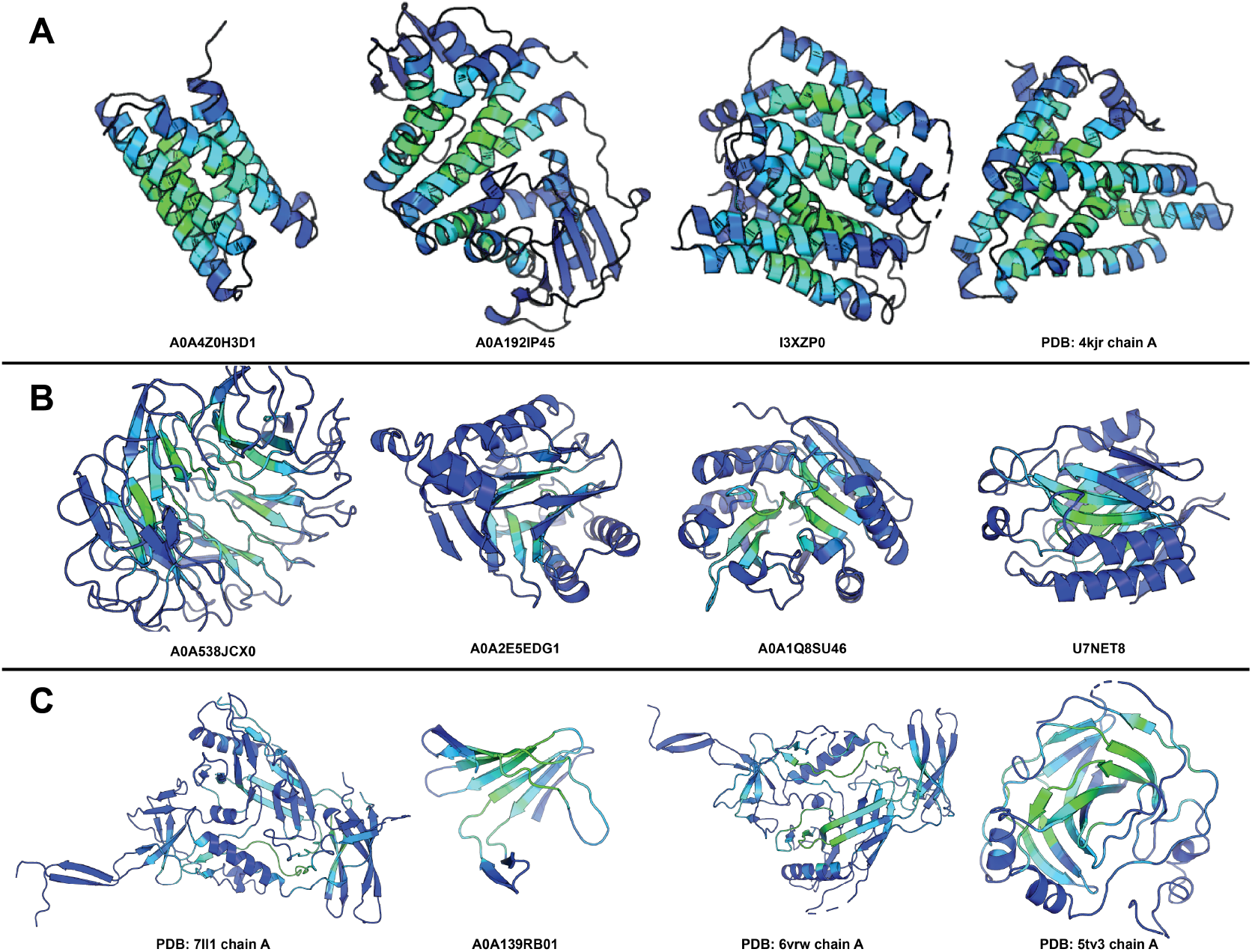
Examples of topics with differing distributions among PDB and AFDB. Topic-specific residue scores are colored from blue (not topic-specific) to green (topic-specific). The difference in frequencies between AFDB (calculated by summing across UniRef50 cluster counts) and PDB chains assigned to each topic is 8% and 3% for **A** and **B** indicating that AFDB has expanded upon these kinds of structural distributions, while it is -3% for **C** indicating that the PDB has proportionally more of this topic. This could be because topic C describes loops near *β*-sheets, while generally loop-filled regions are not high confidence in AlphaFold predictions and hence their shape-mers are not considered.

**Figure 8:**
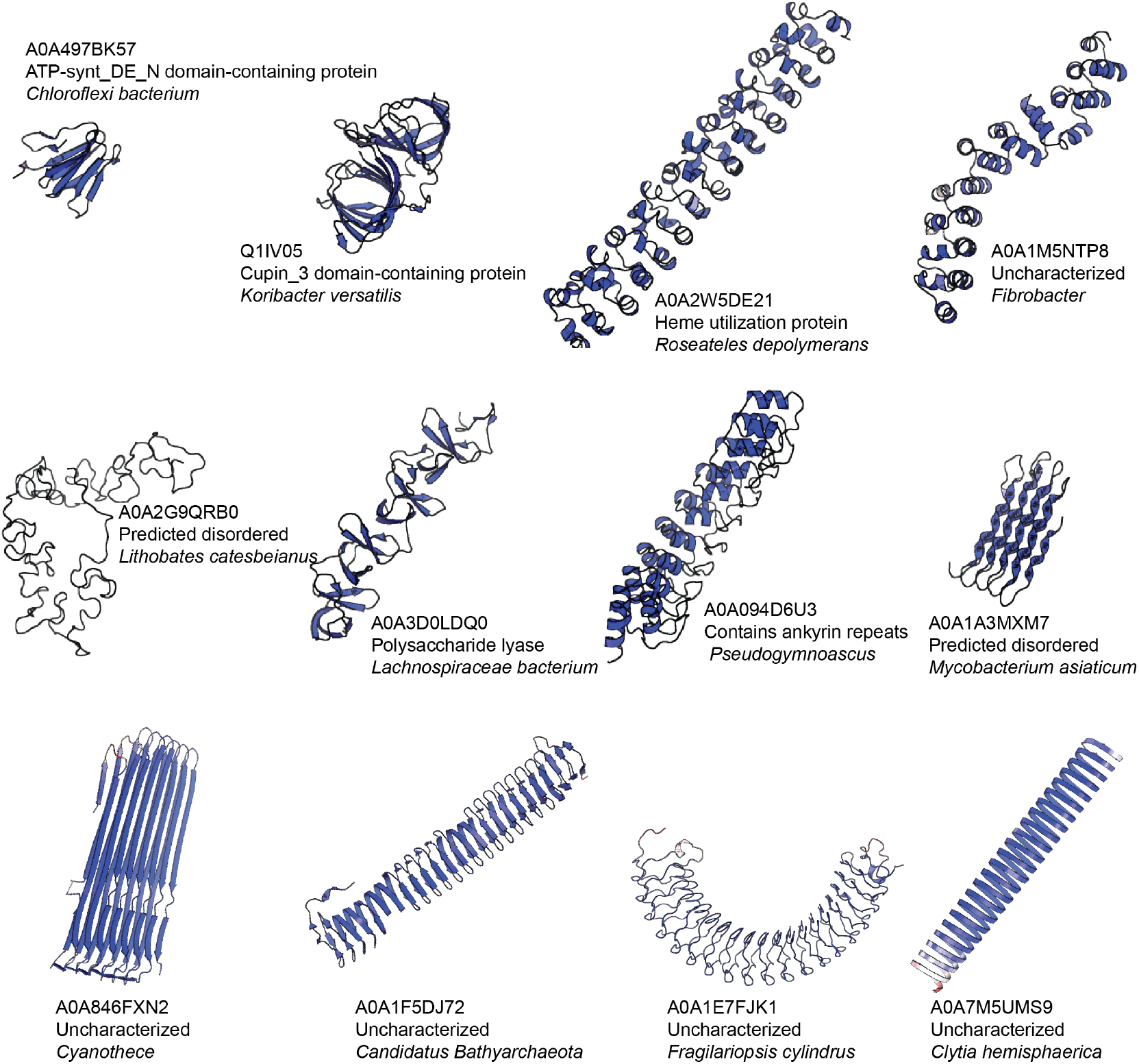
Examples of structural outliers in AFDB with no similar PDB structures (as measured by Word2Vec embedding similarity).

